# The potential role of volatile organic compounds on the colonisation of deadwood by saproxylic beetles

**DOI:** 10.1101/2025.08.29.673086

**Authors:** Claudio Sbaraglia, Simon Thorn, Lucie Ambrozova, Lukas Cizek, Petr Kozel, Daniel Sebastián Rodríguez-León, Thomas Schmitt, Lukas Drag

## Abstract

Volatile organic compounds (VOCs) emitted by deadwood are increasingly recognised as key olfactory cues used by saproxylic beetles to locate suitable substrates. However, how beetle communities respond to these cues during the colonisation of deadwood remains poorly understood. To address this gap, it is essential to quantify both VOC emissions and beetle assemblages, and to experimentally disentangle the main ecological drivers of the deadwood volatilome (i.e., tree species, sun exposure).

To explore the potential role of VOCs on saproxylic beetle colonisation, we exposed 1200 freshly cut branches of oak, beech, spruce, and pine across Central Europe. To mimic natural variation in deadwood and disturbance processes, the bundles were experimentally manipulated either by sterilisation (to reduce endogenous fungi), fungal inoculation with a brown rot fungus (*Fomitopsis pinicola*), a white rot fungus (*Fomes fomentarius*), or burning. From each deadwood bundle, we sampled 448 substances, 89 of which were identified as VOCs and reared 134 saproxylic beetle species.

We observed distinct VOC profiles emitted by broadleaf and conifer species, aligning with the beetles’ tree-type preferences. In conifers, bark beetles, longhorn beetles, and jewel beetles were associated with different chemical cues, whereas beetle taxonomic separation was not observed in broadleaf species. Although experimental treatments altered VOC composition, they could not explain beetle colonisation patterns.

Our study highlights that VOCs emitted during the early decay stage of deadwood are associated with distinct saproxylic beetle assemblages. VOC composition varied across tree species and treatments, suggesting that chemical variation reflects multiple ecological factors. These findings indicate that deadwood diversity, particularly in tree species, contributes to chemical heterogeneity, which is linked to broader beetle assemblages. Forest conservation efforts may therefore need to consider the role of chemical variation in deadwood, as it could influence colonisation patterns of saproxylic beetles and affect the success of biodiversity management strategies.

## 1. Introduction

Deadwood serves as a crucial habitat for numerous groups of organisms that drive its decomposition (Grove, 2002), thereby contributing to nutrient cycling and soil development (Albisetti et al., 2003; Pichler et al., 2012). While fungi and bacteria are the primary decomposers of deadwood (Fukasawa, 2021; Seibold et al., 2022), insects facilitate decomposition both by tunnelling through wood, enhancing microbial colonisation, and by introducing symbiotic fungi and bacteria (Birkemoe et al., 2018). As deadwood decays, there are various changes in its texture and structure, including an increased occurrence of fungi (Ulyshen & Hanula, 2010; Seibold et al., 2023). This leads to variation in the emission of chemical cues, which further influences the composition of deadwood-dependent organisms (Leather et al., 2014).

In deadwood, the composition of Volatile Organic Compounds (VOCs), low-molecular-weight chemical substances that easily evaporate under natural conditions, changes over time, and it is shaped by many factors, such as tree species (Mäki et al., 2021), rot type (Mali et al., 2019), or fungal development stage (Fäldt et al., 1999). Typically, VOCs emitted by fungi during wood degradation are monoterpenes, sesquiterpenes, and oxygenated VOCs, i.e. alcohols, ketones, and acetates with chain-length between four and 15 C-atoms (Evans et al., 2008; El Ariebi et al., 2016). Due to their different wood degradation mechanisms, brown rot fungi tend to emit higher levels of monoterpenes, while white rot species tend to release more protein-based volatiles (Mali et al., 2019). VOC emissions also may vary between fungal structures, with mycelium and sporocarps producing different blends (Orban et al., 2023), which may attract specific species adapted to various ecological niches (Johansson et al., 2006). Because fungi are among the first colonisers of deadwood, their early establishment can lead to priority effects (Chase, 2003). That is, the timing and order of species arrival shape the assembly of later-arriving organisms (Fukami et al., 2010). Early fungal colonisers involved in wood degradation alter VOC emissions (Mäki et al., 2021), potentially influencing the colonisation of saproxylic beetle assemblages attracted to these VOCs.

Saproxylic beetles are a diverse and ecologically important group of organisms associated with deadwood. Their colonisation of deadwood is mediated by a mix of visual and olfactory cues (Jonsell & Nordlander, 1995; Cavaletto et al., 2020). The hypothesis that a single or a few specific compounds drive host localisation in beetles has limited supporting evidence (Fraenkel, 1959; Nottingham et al., 1991), whereas the idea that blends or whole groups of volatiles aid host localisation has received more convincing support in the last decades (Visser & Avé, 1978; Fraser et al., 2003; Bruce et al., 2005). For instance, the synergy between terpenoids (e.g., monoterpenes, sesquiterpenes, diterpenes) emitted by various tree species and the pheromones produced by beetles act as semiochemicals, inducing attraction or repulsion in several bark and wood-boring beetles (Allison et al., 2004; Miller, 2006; Vuts et al., 2016).

The complex interactions among deadwood, wood-inhabiting fungi, and saproxylic beetles potentially mediated by VOCs in early-decay stages remain largely unexplored, despite these stages being widespread and representing key time windows for the colonisation of diverse beetle communities. While most field studies have focused on individual saproxylic beetle species and VOCs from live trees (Seybold et al., 2006; Isidorov et al., 2010; Hulcr et al., 2011), studies focusing on VOCs emitted by deadwood have been mainly conducted under laboratory conditions (Mali et al., 2019; Mäki et al., 2021). As a result, little is known about how beetle communities are associated with VOCs released from fresh deadwood under natural conditions.

To investigate the colonisation process of saproxylic beetles mediated by VOCs, we first collected VOCs from fresh deadwood bundles exposed in natural conditions, and then we sampled the beetle assemblages reared from the same bundles. To simulate natural heterogeneity in deadwood, we created a diverse range of conditions by exposing bundles of different treatments (fungi-depleted, fungi-inoculated, burned) in combination with four tree species (oak, beech, spruce, and pine). This experimentally manipulated approach with a balanced design enabled us to evaluate beetle colonisation choices as a potential response to chemical cues and to test the following hypotheses:

1. VOC composition from deadwood bundles differs among tree species, with conifer and broadleaf species emitting distinct VOC profiles.
2. VOC composition differs among treatments: fungi-inoculated bundles emit distinct VOCs due to differences in decay type (brown vs. white rot), while burned bundles show unique profiles due to thermal degradation.
3. All saproxylic beetles are positively associated with their host tree-specific VOCs; and niche specialists, mycetophagous and pyrophilous beetle species, are positively associated with VOCs emitted by fungi-inoculated and burned bundles, respectively.

## 2. Methods

### 2.1 Study sites

The 10 study sites span a 650 km longitudinal gradient in Central Europe, ranging from 9.68° E to 17.99° E, with elevations between 344 m and 1069 m (Fig. 1a). These sites include diverse forest ecosystems, such as mixed mountain forests (spruce, fir, and beech), lowland oak and pine forests, and upland forests dominated by beech, hornbeam, and maple. (Full site descriptions, including detailed environmental characteristics, are available in (Appendix S1: Table S1)

**Fig. 1.**
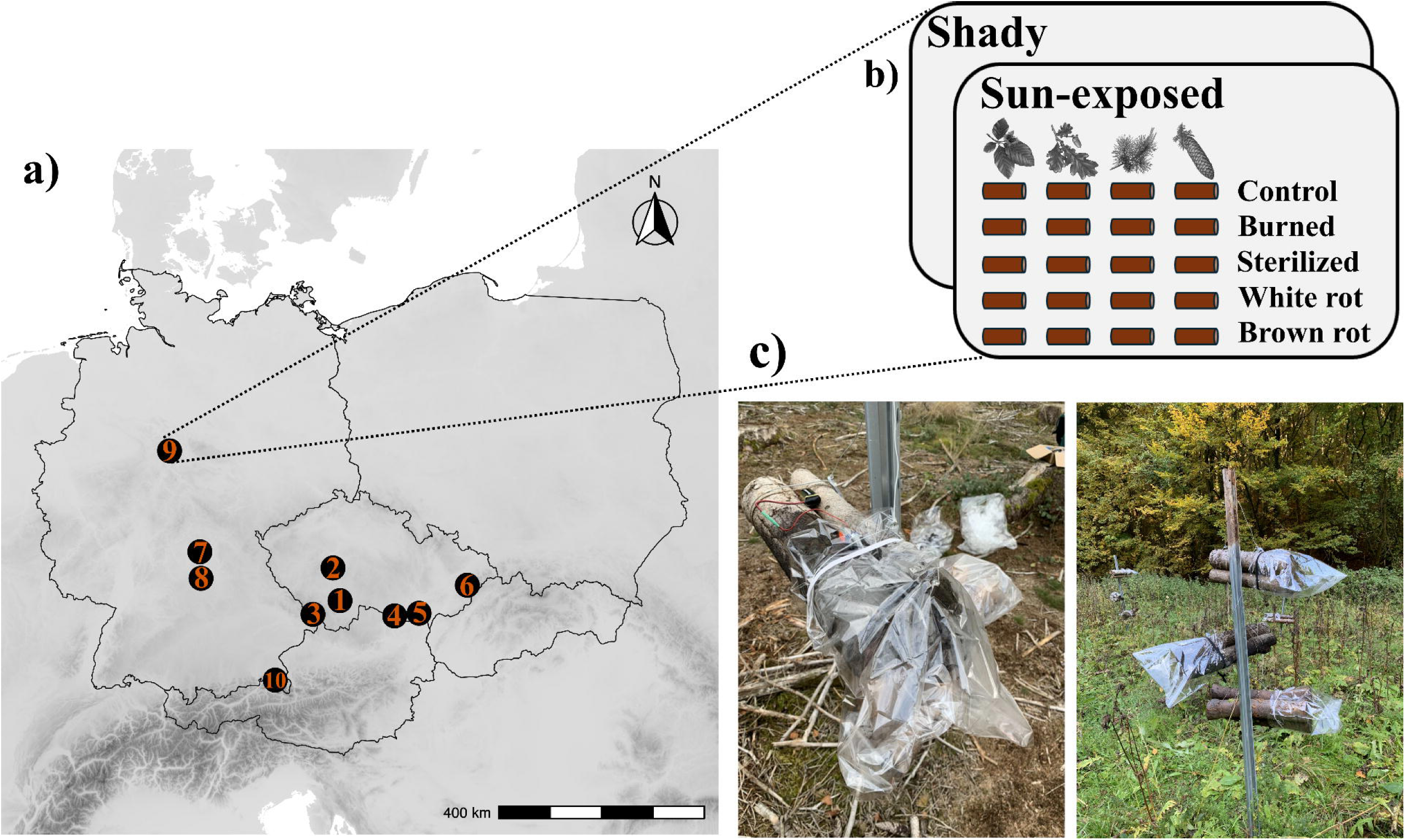
Ten study sites distributed across Germany and the Czech Republic (a). Each site consisted of two plots (shady, sun-exposed), each with 20 bundles (combinations of four tree species and five treatments) (b). VOCs were collected from each bundle during July 2022 after the bundles had been wrapped for 45 minutes to concentrate the volatiles, followed by a 15-minute collection period (c).

### 2.2 Wood collection and processing

Freshly cut logs were obtained in April 2022 from forest sites within a 20 km radius of Markt Erlbach, northern Bavaria (Germany). Each log had a similar size with a mean diameter of 7.8 cm (SD ± 1.33) and a mean length of 51.8 cm (SD ± 0.73). Logs consisted of four tree species: European beech (*Fagus sylvatica*), oak (*Quercus robur/petraea*), pine (*Pinus sylvestris*), and Norway spruce (*Picea abies*). In total, we prepared 1200 logs (300 logs for each tree species). To simulate a broad range of conditions, we treated the bundles by light surface burning, fungal depletion via autoclaving (sterilised), and fungal inoculation with *Fomitopsis pinicola* (brown rot) and *Fomes fomentarius* (white rot) (Appendix S1: Section 1).

### 2.3 Experimental design

We established two plots in each site, one in the forest interior shaded by the forest canopy (hereinafter, as “shaded”) and the other in a nearby open landscape exposed to direct sun for most of the daytime (hereinafter, as “sun-exposed”) (Fig. 1b). A total of 1200 logs were grouped in sets of three, forming 400 bundles, ensuring that logs of the same tree species and treatments were bundled together. Our experimental design thus consisted of 20 bundle combinations (4 tree species * 5 treatments, incl. control) exposed in both shaded and sun-exposed plots, resulting in 40 bundles per site. Two or three bundles were suspended on 2-metre-high iron poles in a random position to minimise the sampling bias, but at the same time avoid the presence of the same treatments or tree species on the same pole. Bundles were exposed from May to October 2022. At the end of the exposure period (October 2022), all 400 bundles from the experimental sites were placed in plastic tubes and stored at the rearing station. Each bundle was placed in a plastic tube with a fine mesh-covered ventilation hole and a collector tube containing propylene glycol as a preservative (Appendix S1: Fig. S1). Reared beetles were collected from the emergence tubes in September 2023 and September 2024 and stored in a freezer until identified to species level. The identified beetles were classified as saproxylic following the criteria in Seibold et al. (2015).

### 2.4 Collection and identification of VOCs

VOC collection was conducted by active headspace sampling using an adsorbent tube and air sampling pump for 15 minutes after the deadwood bundle was enclosed in a polyacetate bag from one side for 45 minutes. The collected samples were analysed using a thermal desorption unit (TDU; TD100-xr, Markes, Offenbach am Main, Germany) connected to a gas chromatograph-mass spectrometer (GC/MS; Agilent 7890B GC and 5977 MS, Agilent Technologies, Palo Alto, USA). We detected 448 peaks in the chromatograms from 400 deadwood bundles across the study sites between July 8–15, 2022, after analysing the volatile traps using gas chromatography/mass spectrometry coupled to the thermal desorption unit.

Because environmental conditions can alter or dilute VOC emissions (Paiva, 2000), all samples were collected under comparable weather conditions, including an average site temperature of ∼25 °C, mostly sunny skies, low wind intensity, and dry bundles (Appendix S1: Table S2), values which suggest a stable and favourable condition for VOCs sampling. Control samples were collected at each site at about 30 metres from the bundles to account for background VOCs. Only peaks emitted at least five times more in bundles than in controls were retained (Appendix S1: Section 2).

### 2.5 Data analysis

All statistical analyses were performed in R (v4.4.3; R Core Team, 2025) through the RStudio platform (RStudio Team, 2024). Figures and data manipulation were carried out using the ‘tidyverse’ package (Wickham et al., 2019).

### 2.6 VOCs selection

To assess how tree species, treatment, and exposure influence VOC emissions, we modelled the relative abundance of each peak (response variable) separately using linear mixed-effect models (LMM) with a Gaussian error distribution from the ‘lme4’ package (Bates et al., 2015). Tree species (oak, beech, spruce, pine), treatment (control, burned, fungi-depleted, brown rot, white rot), and exposure (sun-exposed, shaded) were included as fixed effects. To account for the hierarchical structure of the experiment, the site affiliation was included as a random effect. Peaks with poor model convergence were excluded from further analysis. A total of 89 peaks were retained in the final matrix and used for further analysis. We associated individual peaks with treatments and tree species using Tukey’s post hoc comparisons from the LMM models using the ‘multcomp’ package (Hothorn et al., 2008).

Comparisons were based on model-estimated t-values, with p-values adjusted for multiple testing. Inoculated logs (brown and white rot) were compared to fungi-depleted logs to isolate the effect of inoculation, as both were sterilised. Fungi-depleted and burned logs were compared to untreated controls. For tree species, we tested for species-level differences; when non-significant (e.g., beech vs oak), compounds were assigned to broader categories (“conifer” or “broadleaf”) and tested using the same criteria (Appendix S2).

### 2.7 Composition analysis

To test whether VOC composition in deadwood bundles differed among tree species and treatments, we first conducted a preliminary Detrended Correspondence Analysis (DCA), which revealed a gradient length of 3.39, indicating that a unimodal ordination method was appropriate (Lepš & Šmilauer, 2003). Based on this result, we performed a Canonical Correspondence Analysis (CCA) from the ‘vegan’ package (Oksanen et al., 2001), using the relative abundance of compounds as the response variable. Tree species, treatment, and their interaction were included as explanatory variables, and exposure time as a covariate. To account for differences among study sites, sites were included as a conditioning variable, effectively removing their influence before testing the effects of the explanatory variables. Significant differences among explanatory variables were tested through Permutational Multivariate Analysis of Variance (PERMANOVA) using the ‘adonis2’ function, with 999 permutations. To account for the hierarchical study design, site affiliations were included as strata for permutation constraints. Heatmaps were generated through the ‘ggplot2’ R package (Wickham, 2016) to visualise potential shifts in VOC composition in response to the applied treatments and their interaction with tree species.

### 2.8 Saproxylic beetle preferences

To identify the specific association of saproxylic beetles with tree species and treatments, for each species we calculated Indicator Values (IndVal) using the ‘multipatt’ function in the ‘indicspecies’ package (Cáceres & Legendre, 2009), which quantifies the species’ preference as a percentage. The IndVal is based on two components: specificity (how restricted a species is to a particular group) and fidelity (how consistently it occurs within that group) (Dufrêne & Legendre, 1997). The significance of the association was tested through 999 permutations.

Species with significant IndVal (p < 0.05) for tree species and/or treatments were retained for further analysis.

### 2.9 Association VOC – Beetles

To explore whether beetle colonisation is associated with VOC profiles of specific tree species and treatments, we performed Non-Metric Multidimensional Scaling (NMDS) via the ‘vegan’ package (Oksanen et al., 2001), with four dimensions (k = 4) based on beetle abundance, using 40 species selected through Indicator Species Analysis. To assess the potential association between beetles and VOCs, we fitted the peaks into the NMDS ordination space using the “envfit” function. A significant association was tested through 999 permutations, accounting for the variation in VOC emissions related to sites by permuting within each site. Only significant peaks (p < 0.05) were plotted.

## 3. Results

We detected 89 VOCs primarily emitted from deadwood bundles across the different treatments and tree species. Of these, most peaks were sesquiterpenes (25%) and monoterpenes (22%), followed by hydrocarbons (14%), alcohols (9%), and unidentified compounds (9%). Less represented classes included aldehydes (5%), fatty acids (4%), ethers, ketones, and lactones (each 2%), while disulfides, esters, and furfural were each represented by a single peak. In total, 41 peaks were tentatively identified at the substance level (Appendix S1: Table S3).

Within two years, we reared 134 species of saproxylic beetles (24,696 individuals) from the bundles, with Curculionidae being the most abundant and species-rich family (13,200 individuals across 25 species). The most abundant species were *Xylosandrus germanus* (5091 individuals), *Ernobius mollis* (3397), *Anthaxia quadripunctata* (3231), *Scolytus intricatus* (2668), and *Taphrorychus bicolor* (1468).

### 3.1 VOC composition

We found significant effects of tree species (expl. var. = 16.85%, pseudo-F = 27.43, p = 0.001), treatment (expl. var. = 1.59%, pseudo-F = 1.94, p = 0.001), their interaction (expl. var. = 3.26%, pseudo-F = 1.32, p = 0.003), and exposure (expl. var. = 0.70%, pseudo-F = 3.42, p = 0.001) on the VOC composition (Fig. 2). CCA revealed that VOC collected from deadwood bundles clustered according to broadleaf and conifer tree species, with a clear separation between spruce and pine bundles, but not by treatment (Fig. 2). Heatmaps showed that the relative abundance of VOC composition varied across tree species in response to the applied treatments (Fig. 3). Among the selected peaks, spruce emitted the fewest VOCs compared to other tree species. In conifer bundles, we observed a shift in the relative VOC composition: monoterpenes were more abundant in control bundles, whereas sesquiterpenes emissions increased in fungi-inoculated bundles. In broadleaf bundles, heatmaps indicated increased sesquiterpene emissions in fungi-depleted and fungi-inoculated treatments compared to controls.

**Fig. 2.**
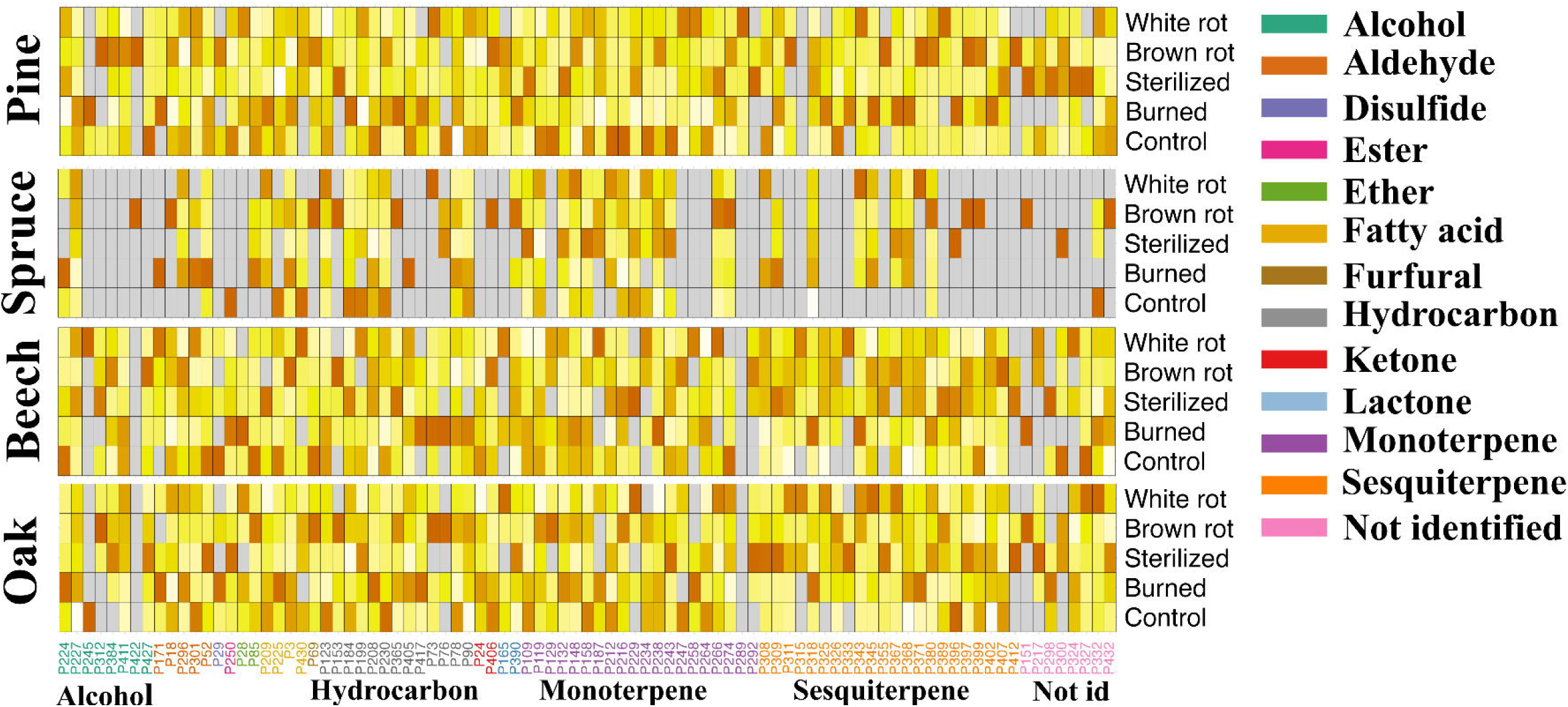
Canonical Correspondence Analysis (CCA) of VOC composition from deadwood bundles. The first two axes explain significantly (*** = p<0.005) 39.5% of the variation. Model-selected peaks (n = 89) were used as response variables; predictors included tree species, treatment, their interaction, and exposure. Bar plots show the variance explained by each predictor. Colours indicate tree species; shapes indicate treatments.

**Fig. 3.**
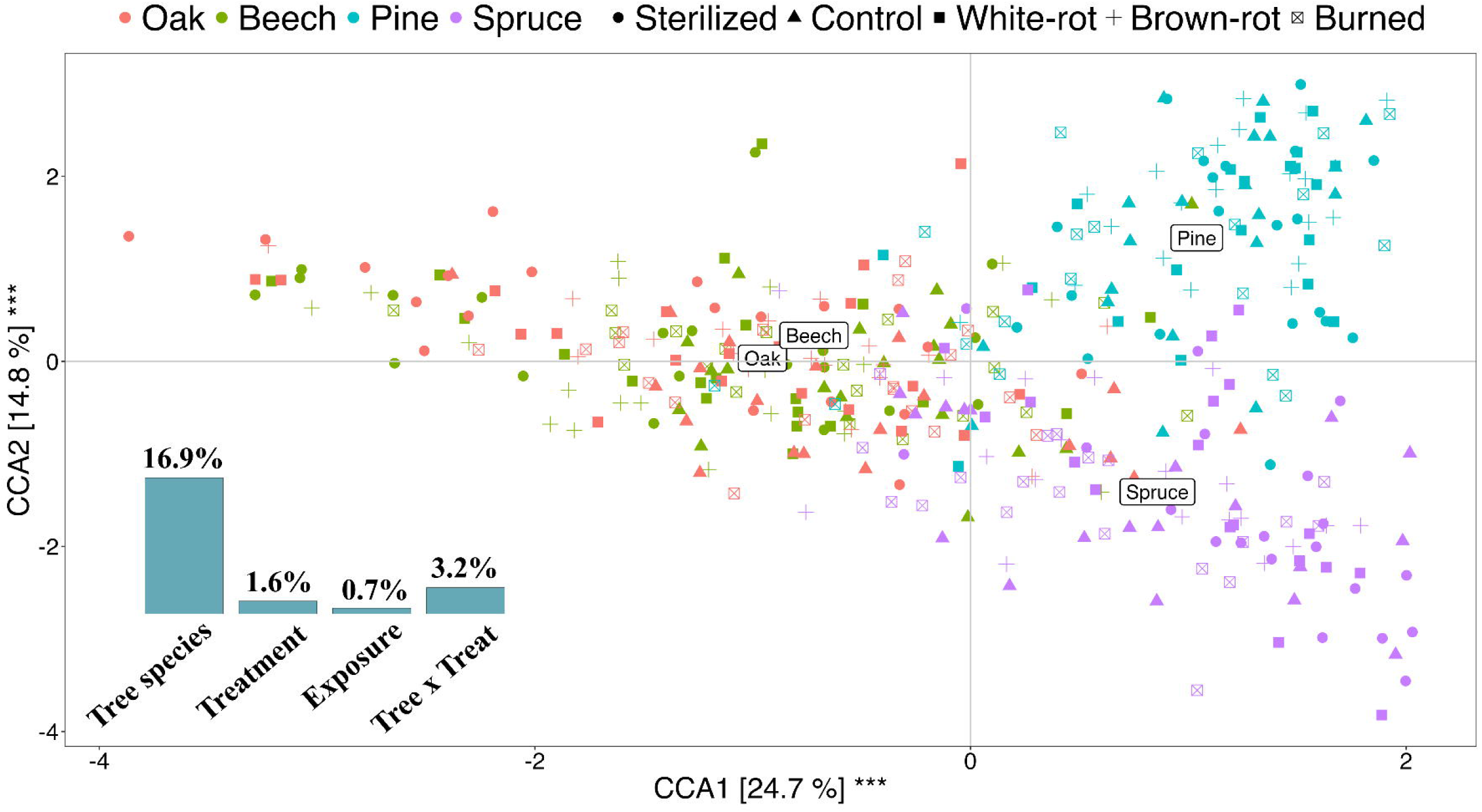
The heatmap showing the average relative VOC composition across tree species and treatment combinations. Peaks are colour-coded by chemical class, with a gradient indicating the level of VOC emissions. Darker colours represent higher emissions, while lighter colours correspond to lower emissions. Grey indicates missing VOCs.

### 3.2 Indicator beetle species

The Indicator Species Analysis identified 45 saproxylic beetle species associated with tree species and 14 associated with treatments (Appendix S3). In our dataset, most beetle species exhibited a preference for a specific tree species. Oak and spruce supported the highest number of beetle species, with 12 and 6 species, respectively. Regarding treatments, *Xylosandrus germanus* and *Synchita mediolanensis* mainly occurred in fungi-depleted and inoculated bundles, while *Leiopus nebulosus* and *Orthoperus atomus* showed a preference for manipulated bundles (fungi-depleted, burned, or fungi-inoculated). Additionally, *Bitoma crenata* and *Cryptophagus scanicus* showed a weak but significant association with brown-rot-inoculated and burned bundles, respectively.

### 3.3 VOCs – Beetle associations

We displayed indicator beetle species and VOC peaks in NMDS ordination space to explore their relationship (Fig. 4). Out of the 42 VOC peaks significantly associated with the ordination axes, 18 showed associations at the species level, seven with pine, five with beech, four with spruce, and two with oak. In cases where differences between species within a tree type (e.g., beech or oak and pine or spruce) were not significant, peaks were broadly categorised by tree type. Based on this grouping, 17 peaks were associated with broadleaf trees and two with conifers. Five peaks did not show any specific tree association. Beetles and VOC peaks clustered by tree species (Fig. 4a), with broadleaf-, beech-, and oak-associated peaks aligning with higher abundances of broadleaf-associated beetles; additionally, peaks characteristic of conifers were associated with increased abundances of spruce-and pine-associated beetles. Regarding treatments, eight peaks were linked to fungi-depleted bundles, two to control, and one each to white rot and burned treatments. None were associated with brown rot, and 30 had no treatment association. Beetles and peaks did not cluster clearly by treatment (Fig. 4b). An exception was the ambrosia beetle *X. germanus*, which was associated with fungi-depleted bundles as well as with some peaks characteristic of this treatment. The burned-associated peak pointed towards *Cryptophagus scanicus*, an indicator for burned bundles.

**Fig. 4.**
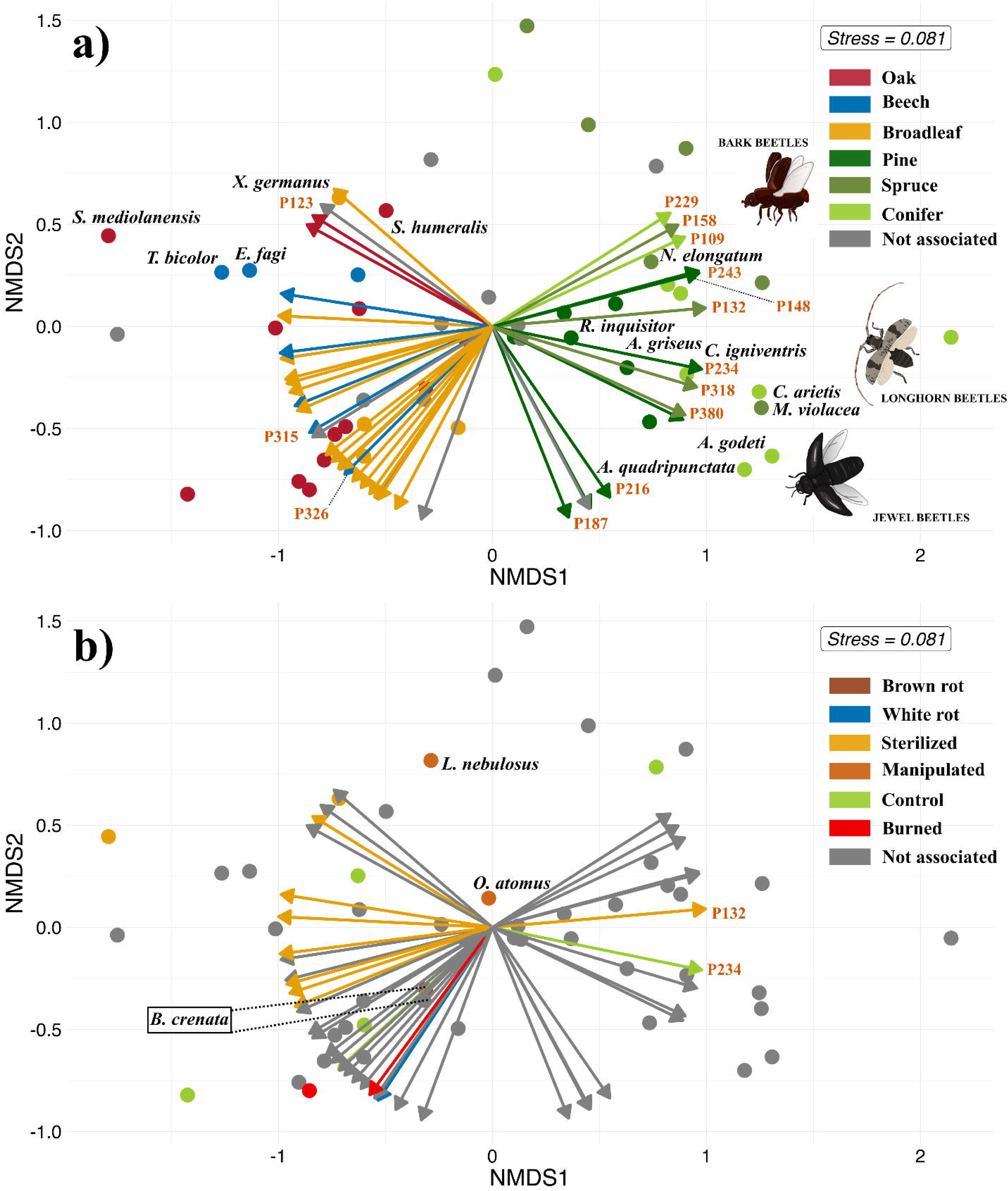
NMDS plots showing associations between VOC peaks and saproxylic beetles by (a) tree species and (b) treatments. Dots represent indicator beetles (n = 40), colour-coded by substrate preference. Grey dots and arrows stand for no preferences from the LMM. Arrows indicate significant VOC peaks (n = 42); their direction indicates positive associations with beetle species (dots) positioned in front of them. Only VOC peaks identified at the substance level are labelled; the full list is available in Appendix S3. Arrow lengths are standardised to unit vectors, not showing effect size (R^2^). The same colour was used for beetles and VOCs associated with the respective tree species (a) or treatment (b). “Manipulated” includes beetles attracted to all treatments except the control. Beetle illustrations by Michaela Helclová.

## 4. Discussion

Our study offers a novel field-based approach combining VOC sampling of deadwood with insect rearing of saproxylic species to investigate chemically mediated colonisation processes in deadwood. We show that VOC composition varies across host tree species and, to a lesser extent, across treatments, and that these chemical differences are associated with distinct saproxylic beetle assemblages. This highlights the potential ecological relevance of host-specific volatile cues during the early stages of deadwood colonisation for saproxylic beetle communities.

### 4.1 Tree species variation in VOCs composition

As hypothesised, tree species accounted for most of the variation in VOC composition (Fig. 2). The distinct chemical profiles between conifers and broadleaves likely reflect differences in wood anatomy and stress-response strategies. Conifers store terpene-rich oleoresin in specialised structures (resin ducts, glands), which release volatiles like monoterpenes and sesquiterpenes upon damage (Franceschi et al., 2005; Raffa, 2014; Celedon & Bohlmann, 2019). In contrast, broadleaf species lack such resin-containing structures and therefore exhibit lower terpene emissions upon physical damage (Holighaus & Schulz, 2006; Thakeow et al., 2007). We observed a distinct VOC composition between spruce and pine bundles (Fig. 2), with pine emitting both a greater number of VOCs and higher levels of monoterpenes in control bundles. These differences likely reflect their contrasting defence strategies: pine relies more on constitutive defences, storing large amounts of resin in preformed ducts, while spruce depends more on induced responses involving de novo synthesis of terpenoids and phenolics (Franceschi et al., 2005; Zhao et al., 2011; Celedón & Bohlmann, 2019). Since VOCs were collected from deadwood, where induced responses are inactive, the stronger emissions from pine likely result from its larger stored resin reserves, whereas spruce’s reliance on inducible pathways may explain its lower VOC emissions.

### 4.2 Fungal colonisation influences VOC composition

The breakdown of wood tissues during fungal colonisation releases a wide array of VOCs (Mäki et al., 2021). In our design, we minimised the a priori fungal presence through sterilisation and used fungi-depleted bundles as controls to isolate the effect of inoculation from sterilisation. Although the variation explained by our manipulated deadwood (inoculation, fungi-depletion, and burning) was low, it significantly influenced VOC composition.

We observed higher relative abundances of monoterpenes in pine control bundles compared to those that were fungi-depleted, inoculated, or burned (Fig. 3). High monoterpene levels in freshly cut logs are expected in conifers, due to the passive release of oleoresin after cutting (Franceschi et al., 2005; Isidorov et al., 2010). Interestingly, this pattern was not observed in spruce bundles, which aligns with findings by Mäki et al. (2021), who reported that monoterpene emissions peak earlier in pine sawdust but later in spruce. However, we found no increase in monoterpene emissions following inoculation in conifers (Fig. 3), suggesting that fungal colonisation did not enhance oleoresin release, possibly due to limited release or rapid degradation of monoterpenes. Alternatively, the heat during the sterilisation and burning process influenced the monoterpenes’ emissions.

Contrary to conifers, beech and oak do not store monoterpenes-rich oleoresin, but rely on tannins and phenolic compounds in their bark as a defence against biotic attacks (Dübeler et al., 1997; Mämmelä, 2001). Fungal degradation of wood in these species is therefore more likely to produce derivatives of these compounds rather than increase monoterpene emissions. In our study, such compounds were indeed detected, including orcinol, thymol, and cymeneol. Their presence, particularly in fungi-inoculated bundles, suggests that fungal activity can enhance the emission of phenolic volatiles, with a stronger effect observed in beech than in oak (Fig. 3).

Sesquiterpene emissions in deadwood are linked to fungal degradation, as observed under laboratory conditions, in pine (Mäki et al., 2021), spruce (Mali et al., 2019), and beech (Holighaus & Schütz, 2006; El Ariebi et al., 2016). In our study, fungi-inoculated bundles in conifers showed higher relative abundances of sesquiterpenes compared to fungi-depleted ones (Fig. 3). However, this pattern was not observed in broadleaf bundles, where differences appeared mainly between control and fungi-depleted bundles. This contrast likely reflects differences in fungal colonisation processes and substrate properties between conifers and broadleaves. In conifer wood, fungal colonisation is generally slower than in broadleaf due to higher lignin content, C:N ratio, and resinous structures (Cornwell et al., 2009). This might explain the relatively low sesquiterpene emissions in fungi-depleted bundles. In contrast, broadleaf wood is more easily colonised due to lower lignin content, allowing environmental fungi to rapidly colonise fungi-depleted bundles. This likely led to similar levels of fungal activity and sesquiterpene emissions in both inoculated and fungi-depleted bundles.

### 4.3 VOCs and beetle responses to burned wood

The burning process alters wood’s structure, texture, and chemical properties, releasing odour cues that are ecologically relevant to pyrophilous beetles across local and landscape scales (Allison et al., 2004; Ramberg et al., 2025). Some of these VOCs (e.g., guaiacol, 5-methylfurfural) act as chemical cues of burned wood for pyrophilous species of longhorn and woodboring beetles. However, on burned bundles, we detected neither pyrophilous beetles nor VOCs typically associated with thermal degradation of cellulose and lignin (Oasmaa et al., 2003; Karagöz et al., 2005), likely due to the two-month delay between the burning event and VOC sampling. Additionally, the amount of burned wood in our experiment was limited and highly fragmented, which may have further reduced its detectability by pyrophilous beetles that rely on long-range chemical cues to locate large-scale fires (Paczkowski et al., 2013) rather than isolated burned logs. In contrast, we detected *Cryptophagus scanicus*, a silken fungus beetle, which associated with burned bundles and an unidentified hydrocarbon (Fig. 4b). This finding suggests a potential case of priority effects, where early fungal colonisers during the degradation of deadwood may have produced volatiles that attracted this species.

### 4.4 VOC beetle association

In the present study, we found that saproxylic beetle communities were associated with VOC profiles of their host trees. Beetle assemblages from spruce and pine occurred in deadwood with higher relative abundances of conifer-associated volatiles (i.e., α-pinene, verbenol), while assemblages from beech and oak were associated with broadleaf-associated volatiles (i.e., sativene, β-copaene) (Fig. 4a). Moreover, the composition of VOC profiles differed markedly between coniferous and broadleaf species: broadleaf bundles emitted a greater diversity of compound classes, predominantly sesquiterpenes, whereas monoterpenes dominated conifer bundles. The distinct separation of beetle communities based on host tree volatiles is particularly striking given the complexity of natural environments, where overlapping odours create a complex mixture of ambient chemical cues. These findings suggest that saproxylic beetles can detect and respond to either a single volatile or blends to locate suitable deadwood for feeding or oviposition.

One distinct cluster of beetles emerging from conifer bundles was dominated by bark beetles and by the predatory species *Nemozoma elongatum*. This cluster was associated with high relative abundances of conifer volatiles such as α-pinene (P109), β-pinene (P132), limonene (P158), verbenone (P243), carene (P148), and terpineol (P229) (Fig. 4a). Several of these monoterpenes are well-known semiochemicals for bark beetles, involved in host location and aggregation (Byers, 1989; López et al., 2013; Sánchez-Osorio et al., 2021), acting as cues. The co-occurrence of *N. elongatum* with bark beetles is likely driven by the presence of its primary prey, *Pityogenes chalcographus*, whose aggregation pheromone, chalcogran, is used by *N. elongatum* as a kairomonal cue for prey detection (Heuer & Vité, 1984).

The second cluster, comprised of jewel beetles, longhorn beetles, and the bark beetle *Magdalis violacea*, linked to cymeneol (P234), longifolene (P318), caryophyllene oxide (P380), and an unidentified compound. Longifolene has been suggested as a potential attractant for *Monochamus galloprovincialis* (Szmigielski et al., 2012), especially when coupled with other common host volatiles (i.e., limonene, α-pinene) (Pajares et al., 2004). In our study, the few individuals detected in a single bundle did not allow us to include *M. galloprovincialis* in our analysis.

Nevertheless, our results suggest that longifolene may serve as a host colonisation cue for other longhorn beetles. For jewel beetles, these associations are reported here for the first time, and further experimental studies are needed to determine whether and how these species detect and respond to these compounds.

A third cluster, including p-cymenene (P187), trans-verbenol (P216), and two unidentified compounds, was not associated with any beetle species in our dataset. Du et al. (2025) reported that under intense bark beetle pressure, emissions of p-cymenene and other aromatic compounds increased in spruce logs before beetle emergence, likely as part of an induced defence response from the logs. This may suggest a deterrent role for these compounds in shaping local beetle assemblages.

Among the broadleaf-associated beetles, the ambrosia beetle *Xylosandrus germanus* was associated with a group of VOCs that comprises 1-ethyl-3-methylbenzene (P123) and three unidentified sesquiterpenes from oak. This ambrosia beetle is known to respond to VOCs emitted by its symbiotic fungus *Ambrosiella grosmanniae* and other fungal sources under bioassay experiments (Mayers et al., 2015; Gugliuzzo et al., 2023). This aligns with our findings, as we recorded thousands of individuals emerging from both fungi-inoculated and fungi-depleted bundles (colonised by pioneer fungi), suggesting that it relies on fungal decay cues to locate and colonise weakened or dead wood in the field.

The second cluster, consisting of a broad range of beetles, including two beech bark beetles, two oak-associated longhorns *Synchita mediolanensis* (Zopheridae) and *Crypturgus dentatus* (Cryptophagidae), and *Litargus connexus* (Mycetophagidae), was linked to a monoterpene emitted by both oak and beech and a sesquiterpene associated with beech.

While their host preferences are consistent with our results, the chemical cues involved in host localisation remain largely unknown. In our study, broadleaf-associated beetles were generally linked to sesquiterpenes (Fig. 4a), a pattern consistent with previous findings (Fäldt et al., 1999; McLeod et al., 2005; Leather et al., 2014). Interestingly, the beech bark beetles, *Taphrorychus bicolor* and *Ernoporicus fagi*, were associated with an unidentified sesquiterpene predominantly emitted from fungi-colonised beech bundles. This association may indicate the use of specific chemical cues to locate early-decay beech substrates.

The third cluster included unidentified sesquiterpenes associated with broadleaf and beech. In our study, this cluster was not linked to any of the beetle species sampled, although these compounds may serve as cues for other species that were either absent or underrepresented in our dataset.

The fourth cluster had a composition similar to the third, consisting of unidentified sesquiterpenes, but was linked to oak-or broadleaf-associated beetles from several families. These compounds may function as general olfactory cues for saproxylic beetles across different families when locating broadleaf deadwood. The only tentatively identified compounds were sativene (P315) and β-copaene (P326), known to be emitted by the brown-rot fungus *Fomitopsis pinicola* (Rösecke et al., 2000), which was used to inoculate a subset of bundles. Although these compounds were more abundant on beech deadwood, we observed a non-significant trend toward higher abundance in brown-rot inoculated bundles compared to white-rot. We suggest that small amounts of these volatiles may serve as host colonisation cues by *Bitoma crenata*, a cylindrical bark beetle associated with brown-rot bundles in our study, suggesting a potential role of priority effects, where early fungal colonisers shape subsequent beetle assemblages through the emission of chemical cues linked to deadwood degradation. Although experimental validation is needed to confirm whether these compounds are detected and behaviourally relevant to beetles, considering aspects like blend composition and concentration, our findings highlight the ecological relevance of VOCs in the colonisation of deadwood by saproxylic beetles.

## 5. CONCLUSIONS

This study is among the first that highlight the role of VOCs in shaping saproxylic beetle communities during the early stage of deadwood decay. We found that beetle assemblages are associated with VOC profiles characteristic of their host tree (conifer or broadleaf), with different groups of volatiles affecting taxa such as bark beetles (Scolytinae), longhorn beetles (Cerambycidae), and jewel beetles (Buprestidae). For some species, the altered VOC profile of fungus-colonised deadwood may have influenced colonisation, suggesting a VOC-mediated priority effect. By focusing on the early stage of decay, our study primarily captured pioneering colonisers, which are typically highly abundant and host-tree specialists, such as bark beetles (Bussler et al., 2011; Seibold et al., 2023). This aligns with our findings of strong host-tree preferences and a dominance of bark beetles. The limited effect of fungal cues on beetle communities may reflect the early successional stage, where chemical cues from host trees play a greater role than those from fungal colonisation, which becomes increasingly influential in later decay stages through fungal–beetle interactions and priority effects.

Although this study focused on beetles, our findings suggest that VOCs may mediate the colonisation processes of other saproxylic organisms (e.g., termites), an aspect worth considering in forest management. Finally, behavioural assays, in the laboratory and the field, are needed to determine whether the identified compounds function as semiochemicals for saproxylic beetles.

## Acknowledgements

This study was supported by the Czech Science Foundation (GACR, project no. 22-27166S). We thank Sven Finnberg for allowing access to part of his forest and for invaluable field assistance. We are also grateful to Jonas Hagge, Max Hetzer, and Michal Perlík for their help during fieldwork, and Lena Unterbauer for managing most of the volatile samples. Special thanks to David Hauck, Jiří Procházka, Pavel Průdek, and František Houška for assistance with beetle identification, and to Michaela Helclová for help with sample sorting and for the beetle illustrations. We appreciate the cooperation and support of Michal Plátek. Thanks to Matthew Sweeney for proofreading the manuscript. We thank the administrations of Podyjí National Park, Pálava PLA, and the Stará Obora game reserve for permitting our research.

## Data Availability Statement

The data that support the findings of this study are available for peer review in the Figshare repository at https://figshare.com/s/680e996049ac87919a90. The dataset will be made publicly available upon publication.

## Author Contributions

**Conceptualization**: Simon Thorn, Lukas Drag, Thomas Schmitt

**Methodology**: Simon Thorn, Lukas Drag

**Formal analysis**: Claudio Sbaraglia, Simon Thorn

**Investigation**: Claudio Sbaraglia, Simon Thorn, Lukas Drag, Lucie Ambrozova, Petr Kozel, Lukas Cizek

**Data curation**: Claudio Sbaraglia, Daniel Rodriguez

**Writing – original draft**: Claudio Sbaraglia

**Writing – review & editing**: Simon Thorn, Lukas Drag, Lucie Ambrozova, Petr Kozel, Daniel Rodriguez, Thomas Schmitt, Lukas Cizek

**Visualization**: Claudio Sbaraglia, Petr Kozel

**Supervision**: Lukas Drag, Simon Thorn, Thomas Schmitt

**Funding acquisition**: Simon Thorn, Lukas Drag

