## Supplementary material for "The potential role of volatile organic compounds on the colonisation of deadwood by saproxylic beetles": Supplementary S1.docx

**Bundle preparation**

Excluding the 240 logs belonging to the control group with no further treatment, the remaining 960 logs underwent one of four treatments: sterilisation (240 logs), sterilisation with the inoculation of brown rot (240 logs) and white rot fungi (240 logs) and burning (240 logs). The number of logs of each tree species was proportional in each treatment category (Fig. 1b). To prevent a priori endophytic fungi contamination, 720 out of 960 logs underwent sterilisation. The logs were enclosed in inert plastic bags and autoclaved for 20 minutes at 120℃. After sterilisation, 480 logs out of the initial 720 underwent drilling at three designated sites (in the middle of the log and on both sides in sections) to allocate dowels covered by fungi mycelium. The dowels had been prepared in the laboratory under controlled conditions, including sterilisation, humidity, and temperature regulation, three months before inoculation into the sterilised logs. This extended preparation period allowed the mycelium to grow sufficiently, maximising the successful colonisation of the logs. Totally, 240 logs were inoculated with brown rot-infested dowels (Red-banded Polypore, *Fomitopsis pinicola,* P. Karst. 1881), and 240 logs with white rot-infested dowels (tinder fungus, *Fomes fomentarius*). Finally, the remaining 240 logs were used for the burning treatment. To simulate the impact of a fire, we manually scorched wood logs for 5 minutes from one side, ensuring a continuum of the burnt surface from unburnt at one end of the log to severely burnt at the other end.

**VOC collection and processing**

VOC collection was conducted by active headspace sampling using an adsorbent tube and air sampling pump. As an adsorbent, we used quartz glass tubes (15 mm × 1.9 mm internal diameter) packed with Tenax (1.5 mg) and Carbotrap (1.5 mg). To keep the adsorbent materials secure, both ends of the tubes were sealed with glass wool, following the method described in Jürgens et al. (2006). Before use, the tubes were conditioned at 250°C for 30 minutes. Each deadwood bundle was enclosed in a polyacetate bag from one side for 45 minutes before the sampling (Otieno et al., 2023). The emitted compounds were drawn through the quartz glass tube for 15 minutes at a flow rate of 1.1 litres per minute using a rotary vane pump (DC12/08FK Fürgut, Tannheim, Germany). The glass tubes containing VOC were separately stored in glass vials in the freezer for GC/MS analysis.

The collected samples were transferred into thermodesorption tubes lined with glass wool, which were loaded into a thermal desorption unit (TDU; TD100-xr, Markes, Offenbach am Main, Germany) connected to a gas chromatograph-mass spectrometer (GC/MS; Agilent 7890B GC and 5977 MS, Agilent Technologies, Palo Alto, USA). The desorption tubes were heated to 260°C for 10 minutes. The desorbed analytes were concentrated in the TDU’s cold trap using nitrogen as the carrier gas and cooled to 5°C. The cold trap was then rapidly heated to 310°C at 60°C per second and maintained for 5 minutes. VOCs were transferred to the GC/MS injector port via a heated transfer line (300°C) using nitrogen. The GC was equipped with an HP-5MS UI capillary column (30 m × 0.25 mm × 0.25 μm, J&W Scientific, Folsom, CA, USA). Helium served as the carrier gas, held at a constant pressure of 1 bar. The GC oven was programmed to start at 40°C, held for 1 minute, then increased to 300°C at 5°C per minute and held for an additional 3 minutes. The transfer line between the GC and MS was maintained at 300°C. The mass spectrometer operated in electron impact (EI) ionisation mode at 70 eV, scanning from m/z 40 to 650 with a scan rate of 2.4 scans per second. To integrate the peaks from the chromatogram, we used Agilent MassHunter Qualitative Analysis Navigator software (v. B.08.00) to process the raw data from our GC/MS analysis.

To efficiently align peaks, we used the ‘GCalignR’ R package (Ottensmann et al., 2018). Peaks eluting beyond 30 minutes were excluded from the analysis, as those with longer retention times are less likely to represent VOCs (Otieno et al., 2023). To remove background emission from the site, peaks were selected only if their mean emission in the bundles was at least five times higher than the mean emission in the corresponding control samples.

VOCs were tentatively identified by comparing their mass spectra to the NIST 2.3 MS Library using Agilent MSD Productivity ChemStation software (MSD ChemStation F.01.03.2357—Agilent Technologies, Inc.). Retention Indices (RI) were calculated from the Retention Times (RT) of n-alkanes, and the values were cross-verified against published RI values and the NIST Chemistry WebBook database. Given the limitations of identification without analytical standards, compounds were primarily classified by their chemical class, with tentative compound names assigned when possible. When classification was not possible, compound classes were labelled as “unknown”. All siloxane-derived compounds were removed from the analysis due to contamination from column bleed (Otieno et al., 2023). When a pair of peaks had closely matching retention indices (RIs), we checked the chromatogram. We compared the peaks to the NIST WebBook to determine whether they represented the same or different compounds and retained them accordingly.

**Figures**

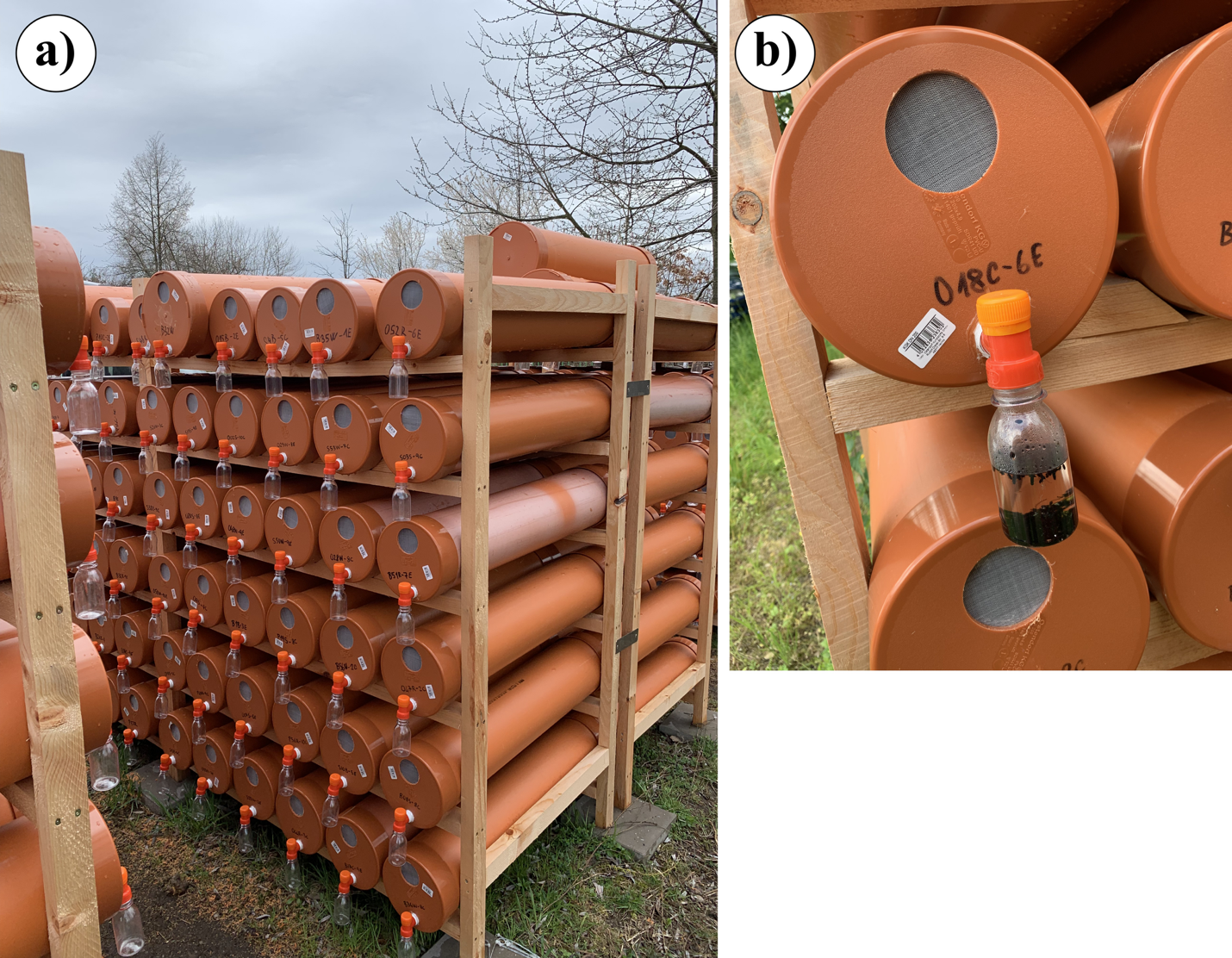

**Fig. S1** Deadwood bundles were stored in plastic tubes with a mesh opening to allow airflow from October 2022 to October 2024 (a), during which emerging beetles were collected in plastic bottles filled with propylene glycol (b).

**Tables**

| **Locality** | **Altitude (masl)** | **Longitude (N)** | **Latitude (E)** | **Distance (m)** | **Land management** | **Forest structure** |
| --- | --- | --- | --- | --- | --- | --- |
| 1 | 538 | 49.09472 | 14.44150 | 75 | Natural monument or feature (III) | game reserve; open mixed forest dominated by *Fagus sylvatica* and *Quercus sp.,* supplemented by *Carpinus betullus*, and *Picea abies*; high amount of deadwood |
| 2 | 390 | 49.66776 | 14.24659 | 244 | Natural area | Open lowland forest dominated by *Pinus sylvestris* with a moderate amount of deadwood |
| 3 | 957 | 48.85445 | 13.68765 | 72 | Protected Landscape (V) | Closed mixed mountain forest with a moderate amount of deadwood |
| 4 | 387 | 48.82706 | 15.96446 | 67 | National Park (II) | Open lowland coppice *Quercus sp.*-dominated forest with a high amount of deadwood |
| 5 | 344 | 48.87260 | 16.64544 | 53 | Habitat/Species Management Area (IV) | Closed *Tilia × europaea* -dominated coppice forest with a low amount of deadwood |
| 6 | 448 | 49.36483 | 17.99042 | 42 | Private land | Shaded forest dominated by *Fagus sylvatica* with a moderate amount of deadwood |
| 7 | 400 | 49.48468 | 10.56609 | 101 | Protected Landscape (V) | Semi-open lowland forest dominated by *Quercus sp.* with a moderate amount of deadwood |
| 8 | 380 | 49.94956 | 10.53332 | 188 | Protected Landscape/ (V) | Semi-open lowland forest dominated by *Quercus sp*. with a moderate amount of deadwood |
| 9 | 410 | 51.71742 | 9.68997 | 116 | Habitat/Species Management Area (IV) | Semi-open lowland forest dominated by *Picea abies* and *Fagus sylvatica* with a moderate amount of deadwood |
| 10 | 1069 | 47.70325 | 12.63522 | 117 | Habitat/Species Management Area (IV) | Closed mixed mountain forest with a moderate amount of deadwood |

**Table S1** Description of the 10 selected sites. For each site, the altitude (m.a.s.l), coordinates (expressed as longitude and latitude), the distance between the shaded plot in the forest interior and the sun-exposed plot outside the forest stand, land management (according to European Environment Agency), and the forest structure describing the canopy openness and amount of deadwood are reported.

| Site | Wind intensity | Cloud cover | Bundles wetness | Temperature (°C) |
| --- | --- | --- | --- | --- |
| 1 | 0 | 1 | 0 | 30 |
| 2 | 0 | 0 | 0 | 30 |
| 3 | 1 | 2 | 1 | 15 |
| 4 | 1 | 0 | 0 | 33 |
| 5 | 1 | 1 | 0 | 34 |
| 6 | 0 | 0 | 1 | 23 |
| 7 | 2 | 1 | 0 | 15 |
| 8 | 1 | 0 | 0 | 25 |
| 9 | 2 | 1 | 0 | 25 |
| 10 | 0 | 2 | 0 | 20 |

**Table S2** Environmental variables were recorded at each study site. Wind intensity (0 = absent/light, 1 = moderate, 2 = strong), cloud cover (0 = clear sky, 1 = partly cloudy with sun, 2 = overcast), bundles wetness (0 = dry, 1 = wet).

| Peak | Compound | Class | RI | NIST (RI) | Tree | Treatment |
| --- | --- | --- | --- | --- | --- | --- |
| P312 | orcinol | alcohol | 1389 | 1370 |  | Brown rot |
| P422 | 4-hydroxy-2-methoxycinnamaldehyde | alcohol | 1739 | 1740 |  | Brown rot |
| P18 | Pentanal | aldehyde |  | 697 |  |  |
| P52 | Hexanal | aldehyde | 802 | 801 |  |  |
| P171 | 2-octenal | aldehyde | 1059 | 1056 |  | Burned |
| P29 | dimethyl disulfide | disulfide |  | 740 |  |  |
| P250 | 2-ethylhexyl acrylate | ester | 1232 | 1224 | Broadleaf |  |
| P28 | 1,1-diethoxy-ethane | ether |  | 727 |  | Burned |
| P85 | n-butylether | ether | 883 | 888 |  | Control |
| P3 | acetic acid | fatty acid |  | 646 |  |  |
| P430 | tetradecanoic acid | fatty acid | 1769 | 1765 |  |  |
| P69 | furfural | furfural | 832 | 834 |  | Brown rot |
| P76 | ethylbenzene | hydrocarbon | 858 | 856 |  |  |
| P90 | styrene | hydrocarbon | 894 | 895 |  |  |
| P123 | 1-ethyl-3-methyl-benzene | hydrocarbon | 962 | 964 |  |  |
| P230 | naphthalene | hydrocarbon | 1183 | 1178 |  | Control |
| P73 | 2,4-dimethyl-1-heptene | hydrocarbon | 842 | 843 | Beech |  |
| P78 | 4-methyl-octane | hydrocarbon | 861 | 864 | Beech |  |
| P165 | lilac lactone | lactone | 1042 | 1041 |  |  |
| P109 | α-pinene | monoterpene | 932 | 939 | Conifer |  |
| P119 | thujadiene | monoterpene | 952 | 956 |  |  |
| P132 | β-pinene | monoterpene | 975 | 978 | Spruce | Sterilised |
| P148 | carene | monoterpene | 1010 | 1007 | Pine |  |
| P158 | limonene | monoterpene | 1029 | 1030 | Spruce |  |
| P187 | p-cymenene | monoterpene | 1090 | 1088 | Pine |  |
| P216 | trans-verbenol | monoterpene | 1146 | 1141 | Pine |  |
| P229 | terpineol | monoterpene | 1178 | 1177 | Conifer |  |
| P234 | cymeneol | monoterpene | 1187 | 1182 | Pine | Control |
| P238 | myrthenol | monoterpene | 1197 | 1196 |  |  |
| P243 | verbenone | monoterpene | 1211 | 1206 | Pine |  |
| P247 | cuminaldehyde | monoterpene | 1224 | 1224 |  |  |
| P274 | thymol | monoterpene | 1296 | 1297 | Oak |  |
| P289 | 2-methyl-5-(propan-2-ylidene)cyclohexane-1,4-diol | monoterpene | 1329 | 1321 | Pine | White rot |
| P315 | sativene | sesquiterpene | 1398 | 1396 | Beech |  |
| P318 | longifolene | sesquiterpene | 1408 | 1402 | Spruce |  |
| P326 | β-copaene | sesquiterpene | 1432 | 1430 | Beech |  |
| P343 | γ-muurolene | sesquiterpene | 1480 | 1474 |  |  |
| P345 | curcurmene | sesquiterpene | 1485 | 1485 |  |  |
| P355 | cuparaene | sesquiterpene | 1509 | 1500 | Broadleaf | Sterilised |
| P367 | α-calacorene | sesquiterpene | 1547 | 1548 |  | White rot |
| P380 | caryophyllene oxid | sesquiterpene | 1587 | 1581 | Spruce |  |

**Table S3** List of volatile organic compounds (n = 41) tentatively identified at the compound level and grouped alphabetically by chemical class. The table includes the calculated Retention Index (RI) and corresponding values from the NIST WebBook, where available. Associations based on IndVal values with tree species and/or treatments (e.g., burned, brown rot) are indicated in the final two columns.
